# Development and Validation of a High-Throughput Short Sequence Typing Scheme for *Serratia marcescens* Pure Culture and Environmental DNA

**DOI:** 10.1101/2021.07.01.450819

**Authors:** Thibault Bourdin, Alizée Monnier, Marie-Ève Benoit, Emilie Bédard, Michèle Prévost, Caroline Quach, Eric Déziel, Philippe Constant

**Affiliations:** Institut national de la recherche scientifique, Centre Armand-Frappier Santé Biotechnologie, 531 boulevard des Prairies, Laval (Québec), Canada, H7V 1B7; Department of Civil, Geological, and Mining Engineering, Polytechnique Montréal, CP 6079, Succ. Centre-ville, Montréal, QC, Canada H3C 3A7; Université de Montréal, Montréal, QC, Canada

**Author notes:** Address correspondence to Eric Déziel, and Philippe Constant. **E-mail address of authors:** T. Bourdin; A. Monnier; M-È. Benoit; E. Bédard; M. Prévost; C. Quach; E. Déziel; P. Constant.

**Keywords:** Molecular typing, HiSST, Sink environment, Opportunistic pathogens, Neonatal intensive care units (NICU), healthcare-associated infections (HAI)

## Abstract

Molecular typing methods are used to characterize the relatedness between bacterial isolates involved in infections. These approaches rely mostly on discrete loci or whole genome sequences (WGS) analyses of pure cultures. On the other hand, their application to environmental DNA profiling to evaluate epidemiological relatedness amongst patients and environments has received less attention. We developed a specific, high-throughput short sequence typing (HiSST) method for the opportunistic human pathogen *Serratia marcescens*. Genes displaying the highest polymorphism were retrieved from the core genome of 60 *S. marcescens* strains. Bioinformatics analyses showed that use of only three loci (within *bssA, gabR* and *dhaM*) distinguished strains with the same level of efficiency than average nucleotide identity scores of whole genomes. This HiSST scheme was applied to an epidemiological survey of *S. marcescens* in a neonatal intensive care unit (NICU). In a first case study, a strain responsible for an outbreak in the NICU was found in a sink drain of this unit, by using HiSST scheme and confirmed by WGS. The HiSST scheme was also applied to environmental DNA extracted from sink-environment samples. Diversity of *S. marcescens* was modest, with 11, 6 and 4 different sequence types (ST) of *gabR, bssA* and *dhaM* loci amongst 19 sink drains, respectively. Epidemiological relationships amongst sinks were inferred on the basis of pairwise comparisons of ST profiles. Further research aimed at relating ST distribution patterns to environmental features encompassing sink location, utilization and microbial diversity is needed to improve the surveillance and management of opportunistic pathogens.

## Introduction

Interactions between patients and the built environment of the hospital has gained attention in epidemiological studies aimed at identifying origins of nosocomial outbreaks. For instance, sink environments are recognized as a source of opportunistic pathogens in several healthcare-associated infections (HAI) (1–3). In preventive or outbreak investigations, molecular typing methods are commonly used to examine the relatedness of environmental or clinical isolates. Many typing techniques are available to achieve this goal (4–10), mostly based on multilocus sequence typing (MLST) methodologies initially developed by Maiden et al. (11). Democratization of high-throughput sequencing technologies have contributed to expand public genome databases, providing an unprecedented portrait of microbial diversity. This has led to the realization that the pangenome of bacterial species displays a mosaic landscape supporting the metabolic flexibility necessary to ensure species resistance and resilience towards disturbances. Such plasticity of microbial genome highlights the need to update and revisit conventional MLST schemes, typically relying on housekeeping genes. In some instances, these genes are not specific enough for accurate molecular typing of investigated strains (12, 13).

The genus *Serratia* is a Gram-negative bacterium classified as members of Enterobacteriaceae that are ubiquitous in water, soil, plants and different hosts including insects, humans and other vertebrates (14, 15). Amongst *Serratia* species, *Serratia marcescens* is the most important opportunistic human pathogen, often multidrug resistant and involved in outbreaks of HAI in neonatal intensive care units (NICU) (16–26). No MLST scheme exists for the molecular typing of *S. marcescens* but other typing techniques have been used during previous epidemiological studies, such as pulsed-field gel electrophoresis (16, 18, 22), ribotyping (27) or more recently whole-genome MLST (28). Even though these techniques were proven efficient to distinguish strains, they are not tailored to epidemiological surveys involving large sample size because they are technically demanding due to upstream cultivation and isolation efforts.

This study introduces a new molecular typing approach, that we called High-Throughput Short Sequence Typing (HiSST), to detect and identify *S. marcescens* relying on culture-dependent and culture-independent applications. The HiSST method was developed based on whole genome sequences of *S. marcescens* available in public databases then validated with reference culture collections, clinical isolates and environmental DNA samples.

## Materials and Methods

### Development of the HiSST scheme

A pan-genome allele database was assembled from 60 complete genomes of *S. marcescens* retrieved from the NCBI GenBank database (last updated in July 2020) with the *Build_PGAdb* module available on PGAdb-builder online tool (29). Conserved genes showing the highest number of alleles were selected as the most variable and discriminant. Alleles of each selected genes (*n* = 32) were aligned, non-overlapping ends were removed and sequence identity matrix was computed with the software BioEdit (30). Gene fragments (< 350 bp) displaying the highest variability were chosen and aligned against the NCBI database with the Basic Local Alignment Search Tool (BLAST) to assess specificity. The 15 loci (i.e. nucleotide sequences of internal fragments of the previously selected genes) showing the highest variability and with the most specific non-overlapping ends were selected as candidates for the HiSST scheme. A trade-off between the number of different loci and specificity of the assay was achieved by topology data analysis of concatenated HiSST loci and genome similarity. Three successive steps were necessary to implement the approach relying on the 60 complete genomes of *S. marcescens* (Table S1) and 9 other strains of non-*marcescens Serratia* spp. available on “GenBank” of NCBI. First, Average Nucleotide Identity (ANIb) (31) analyses were performed on the 69 complete genomes with BLAST+ alignment tool (https://ftp.ncbi.nlm.nih.gov/blast/executables/blast+/LATEST/) with Python package *pyani* (https://github.com/widdowquinn/pyani) (32). Second, a stepwise approach was implemented to assemble concatenated loci alignments. Backward selection procedure was applied on the 15 most discriminant loci until only three loci remained. This led to multiple concatenated alignments comprising either 15, 7, 4 or 3 gene fragments. ANIb scores were calculated on the concatenated alignments of loci. Third, the discriminating power of the four different HiSST schemes was validated by topology data analysis of concatenated loci and complete genomes trees. The topology of the different UPGMA dendrograms were compared with R version 4.0.4 (33) using the packages *pvclust* (34), *dendextend* (35) and *tidyverse* (36). Further, ANIb values of the selected HiSST loci and complete genomes were compared using the packages *circlize* (37) and *ComplexHeatmap* (38), which help to visualize the difference of the discriminatory power between HiSST scheme and whole genome of *S. marcescens*. The R scripts we developed are available on GitHub (https://github.com/TBourd/R_scripts_for_HiSST_scheme). Following these analyses, the three gene fragments of *gabR* (HTH-type transcriptional regulatory protein), *bssA* (Benzylsuccinate synthase alpha subunit) and *dhaM* (PTS-dependent dihydroxyacetone kinase, phosphotransferase subunit) were retained for the further development of the HiSST scheme, as described below.

### Primer design and PCR amplification of *gabR, bssA* and *dhaM* internal loci

Oligonucleotides comprising 18 to 22-mers with either a single or no substituted base were designed to target discriminant internal loci in the *gabR, bssA* and *dhaM* genes (Table 1). *In-silico* tests of primers were performed with the software tool “Primer-BLAST” (39), using the RefSeq non-redundant proteins database to assess specificity and the *Serratia marcescens* subset RefSeq database to verify the coverage of primers for this species. The reaction was carried out in 25 µL of master mix containing 2.5 U/µL Fast-Taq DNA polymerase (Bio Basic Inc., Markham, Canada), 1x of Fast-Taq Buffer (Bio Basic Inc., Markham, Canada), 200 µM dNTPs, 0.4 mg/mL BSA (Bovine Serum Albumin), 0.4 µM of each primer, and 2 ng/µL of extracted DNA. A solution of 0.5x Band Sharpener (Bio Basic Inc., Markham, Canada) was included for the *gabR* mixture only. PCR conditions were optimised for each primer sets with genomic DNA of *S. marcescens* strains as template (Table 1).

**Table 1:**
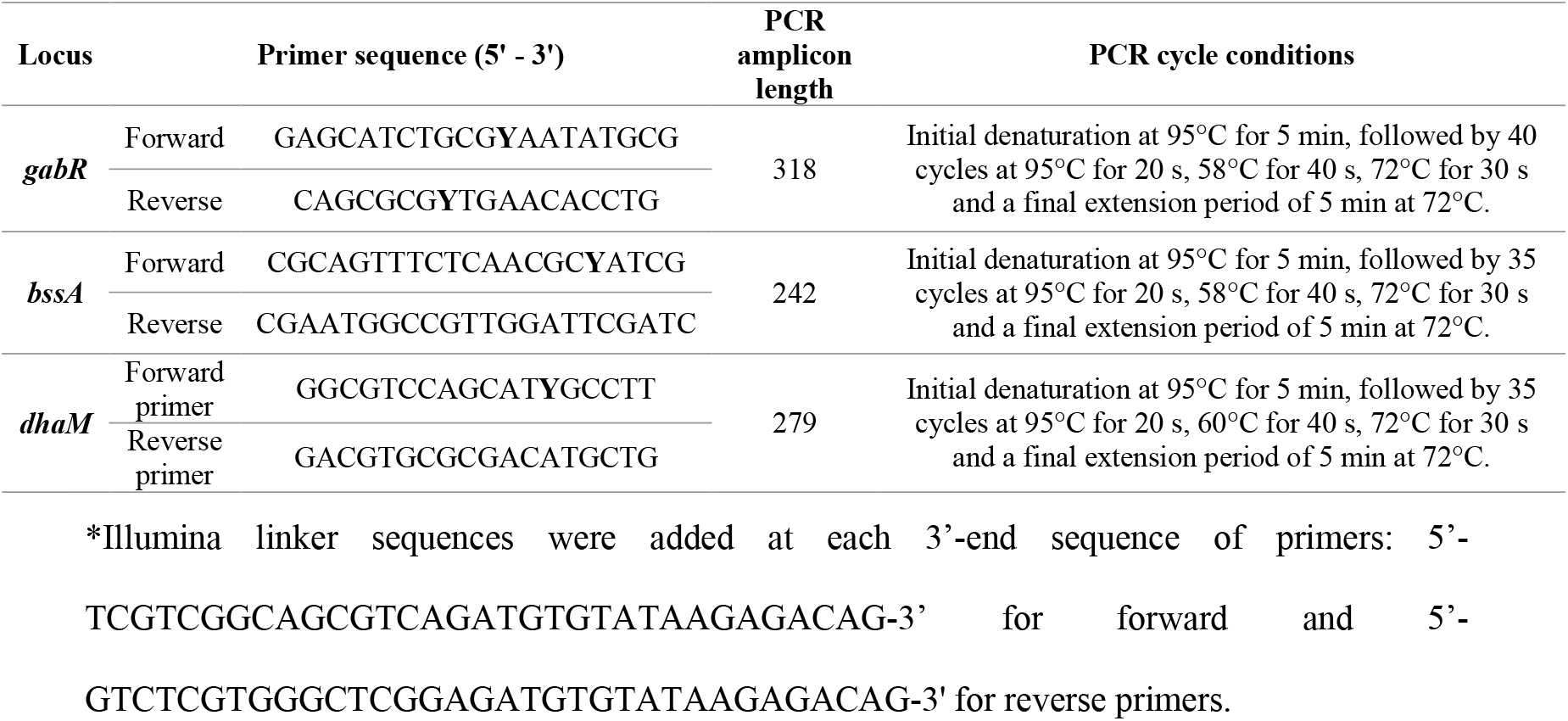
HiSST locus specific primers sequences and PCR cycle conditions^*^.

### Validation of the HiSST scheme with reference strains

Validation of primers was done with 28 reference strains various origins (Table 2). Selected strains comprised *S. marcescens* (*n* = 15), *Serratia rubidaea* (*n* =1), *Serratia liquefaciens* (*n* = 1), *Serratia plymuthica* (*n* = 1), *Pseudomonas aeruginosa* (*n* = 3), *Klebsiella pneumoniae* (*n* = 1), *Stenotrophomonas maltophilia* (*n* = 4), *Stenotrophomonas acidaminiphila* (*n* = 1) and *Stenotrophomonas nitritireducens* (*n* = 1). The strains were purified on Trypticase Soy Broth (TSB) (Difco Laboratories, Sparks, MD, USA - Le pont de Claix, France) with Agar (15 g/L) (Alpha Biosciences, Inc., Baltimore, MD, USA) at 30°C for 48 h. A single colony of each strain was inoculated in 2 mL TSB and grown for 48 h at 30°C for subsequent genomic DNA extraction.

**Table 2.** Reference strains utilized as positive or negative control for HiSST scheme validation. Ten strains of *Serratia* spp. were included to verify the specificity for S. *marcescens* of each selected locus, with *S. ficaria* (*n* = 1), *S. liquefaciens* (*n* = 5), *S. quinivorans* (*n* = 2), *S. proteamaculans* (*n* = 2), also downloaded from NCBI Genome database. These *Serratia* spp. have the most similar nucleotide sequences of selected locus according to the results of BLAST run.

| Species and strains<br>(additional designation) | Lab<br>collection<br># | Isolation origin (country) | Provided by |
| --- | --- | --- | --- |
| *Serratia marcescens* ($n = 15$) | | | |
| PCI 1107 (ATCC 14756, LMG 13576) | ED3691 | Fort Detrick, Maryland (USA) | BCCM-LMG<sup>c</sup> |
| BD1b-2wD | ED4305 | Drain water from a NICU (Canada) | Lab strain |
| Db11 | ED3837 | Insect isolate, *Drosophila melanogaster* (France) | A. Brassinga<sup>a</sup> ; J. J. Ewbank<sup>b</sup> |
| Db10 | ED3838 | Insect isolate, *Drosophila melanogaster* (France) | A. Brassinga<sup>a</sup> ; J. J. Ewbank<sup>b</sup> |
| BS 303 (ATCC 13880, LMG 2792) | ED3696 | Pond water (Czech Republic) | BCCM-LMG<sup>c</sup> |
| L00128734 | ED3957 | Human clinical specimen (Canada) | LSPQ<sup>d</sup> |
| L00128736 | ED3958 | Human clinical specimen (Canada) | LSPQ<sup>d</sup> |
| L00128737 | ED3959 | Human clinical specimen (Canada) | LSPQ<sup>d</sup> |
| L00128966 | ED3960 | Human clinical specimen (Canada) | LSPQ<sup>d</sup> |
| L00128967 | ED3961 | Human clinical specimen (Canada) | LSPQ<sup>d</sup> |
| L00129585 | ED3962 | Human clinical specimen (Canada) | LSPQ<sup>d</sup> |
| L00130169 | ED3963 | Human clinical specimen (Canada) | LSPQ<sup>d</sup> |
| L00133794 | ED3964 | Human clinical specimen (Canada) | LSPQ<sup>d</sup> |
| L00134617 | ED3965 | Human clinical specimen (Canada) | LSPQ<sup>d</sup> |
| L00085643 | ED3966 | Environmental (Canada) | LSPQ<sup>d</sup> |
| *Serratia rubidaea* ($n = 1$) | | | |
| FB299 | ED3693 | Environmental (USA) | Bernier *et al.*, 1994 |
| *Serratia liquefaciens* (n = 1)<br>ID150497 | ED3967 | Human clinical specimen<br>(Canada) | LSPQ<sup>d</sup> |
| *Serratia plymuthica* (n = 1)<br>ID157970 | ED3968 | Human clinical specimen<br>(Canada) | LSPQ<sup>d</sup> |
| *Stenotrophomonas maltophilia* (n = 4)<br>560 (ATCC 13636, LMG 961, NCTC<br>10258) | ED3699 | Human, cerebrospinal fluid<br>(USA) | BCCM-LMG<sup>c</sup> |
| L00083595 | ED3969 | Human clinical specimen<br>(Canada) | LSPQ<sup>d</sup> |
| L00092250 | ED3970 | Human clinical specimen<br>(Canada) | LSPQ<sup>d</sup> |
| L00124341 | ED3971 | Human clinical specimen<br>(Canada) | LSPQ<sup>d</sup> |
| *Stenotrophomonas acidaminiphila* (n = 1)<br>L00129488 | ED3979 | Human clinical specimen<br>(Canada) | LSPQ<sup>d</sup> |
| *Stenotrophomonas nitritireducens* (n = 1)<br>ATCC BAA-12 (LMG 22074, DSM<br>12575) | ED3701 | Laboratory scale biofilter<br>(Germany) | BCCM-LMG<sup>c</sup> |
| *Pseudomonas aeruginosa* (n = 3)<br>UCBPP-PA14 | ED1 | Human clinical specimen<br>(USA) | Daniel G Lee *et al.*,<br>2006 |
| PAO1 | ED956 | Human, wound (Australia) | Sylvie Chevalier<sup>e</sup> |
| FKS4A | ED0129 | Human, cystic fibrosis (USA) | Luke Hoffman<sup>f</sup> |
| *Klebsiella pneumoniae* (n = 1)<br>ATCC 4352 (LMG 3128) | ED3692 | Cow's milk | ATCC<sup>g</sup> |

<sup>a</sup>Ann Brassinga, University of Manitoba, Winnipeg, MB, Canada.
<sup>b</sup>Jonathan J. Ewbank, University of Aix-Marseille, Marseille, France.
<sup>c</sup>Belgian Coordinated Collections of Microorganisms, University of Gent, Belgium.
<sup>d</sup>Laboratoire de Santé Publique du Québec, Sainte-Anne-de-Bellevue, QC, Canada.
<sup>e</sup>Sylvie Chevalier, University of Rouen-Normandie, Rouen, France.
<sup>f</sup>Luke Hoffman, Seattle Children's, Seattle, WA, USA.
<sup>g</sup>American Type Culture Collection, Rockville, MD, USA.

### Validation of the HiSST scheme with environmental DNA

Biofilm and 50 mL of water from ten different sink drains were sampled on April 2019 during an outbreak of *S. marcescens* in a neonatal intensive care unit (NICU) in a Montreal Hospital (Québec, Canada). The same day, samples were inoculated on a semi-selective DNase test agar (40) supplemented with ampicillin (5 μg/ml), colistin (5 μg/ml), cephalothin (10 μg/ml), and amphotericin B (2.5 μg/ml) incubated for 48 h at 30°C. Colonies were purified on TSB with Agar at 30°C for 48 h. During a second sampling campaign, biofilm and water (50 mL) from sink drains were collected twice from 19 sinks in January 2020. Samples were kept on ice during their transportation to the laboratory. Genomic DNA from isolated strains and environmental samples was extracted by a procedure combining mechanical and chemical lysis, using bead beater and ammonium acetate treatment, as previously described (41), prior to PCR amplicon sequencing. The two successive PCR amplifications necessary for the preparation of *gabR, bssA* and *dhaM* sequencing libraries were conducted with the AccuPrime™ *Taq* DNA Polymerase System, High Fidelity (Invitrogen Ltd, Carlsbad, USA). PCR conditions and reaction mixtures were adapted following manufacturer instructions (Table 1). The first PCR reaction was performed using modified *gabR, bssA* and *dhaM* primers including Illumina linker sequences (Table 1) and 2 ng/µL of template DNA. PCR products were purified with AMPure XP beads (Beckman Coulter Inc., Brea, USA). Purified PCR products were subjected to a second PCR performed for libraries preparation using barcoded primers (Table S2) supplied by Integrated DNA Technologies Inc. (Mississauga, Canada). Purified PCR amplicons were quantified using the Quant-iT™ PicoGreen™ dsDNA Assay Kit (Invitrogen Ltd, Carlsbad, USA), diluted and pooled together into 75 µL comprising 1.5 ng/μL of DNA final concentration before shipping for sequencing. PCR amplicons were sequenced with the Illumina MiSeq PE-250 platform at the Centre d’expertise et de services Génome Québec (Montréal, Canada). Raw sequencing reads processing included primer sequences removal with the software Cutadapt v. 2.10 (42), followed by quality control, paired ends merging and chimera check using the default parameters specified in the package dada2 v1.8.0 (43) that include packages ShortRead v1.48.0 (44) and Biostrings v2.58.0 (45). Reads containing a mismatch in the primer region were deleted (R script available on https://github.com/TBourd/R_scripts_for_HiSST_scheme). Filtered sequences were clustered into amplicon sequence variants (ASV) displaying 100% identity. ST assignation of chimera-free ASV was done using *gabR, bssA* and *dhaM* reference databases with a 100% identity cutoff (Table S1). Proportion of reads remaining after each step of the bioinformatics pipeline is provided in Table S2.

### SNP and HiSST-profile analyses

SNPs of each locus were analyzed from all unique nucleotide sequence for references strains and environmental DNA (Table S3). For each strain, the combination of alleles at each locus defined the sequence type (ST), and the combination of multilocus ST defined the HiSST-profile. SNP matrix and HiSST-profile were analysed by geoBURST algorithm using PHYLOViZ platform, version 2.0a (46), creating minimum spanning trees using default software settings. The coverage of the HiSST scheme was visualized in a chart representing the cumulative frequency of ST depending on the number of cumulative loci, based on 60 references strains of *S. marcescens* (Table S1).

### Validation of molecular typing by whole genome sequencing (WGS)

Environmental strain BD1b-2wD and clinical strains ED3957, ED3958, ED3959 were subjected to whole genome sequencing (WGS) with the Illumina NextSeq 550 platform at the Microbial Genome Sequencing Center (Pittsburgh, PA, USA). A quality control of the WGS data was checked with FastQC tool (https://www.bioinformatics.babraham.ac.uk/projects/fastqc). Illumina adapter clipping and quality trimming were performed with Trimmomatic v0.39 (47), by specifying an average quality required greater than 30. Then, genomes were assembled from fastq file of paired-end reads using SPAdes *de novo* assembler (48), and Bandage (49) to visualize SPAdes graphical output. Contigs obtained from SPAdes output were aligned, ordered and oriented with the closest reference genome (*S. marcescens* AR_0122) to create a contiguated genome by using ABACAS tool (50). Pairwise comparison of contiguated genomes of DB1b-2wD, ED3957, ED3958 and ED3959 was performed using ANIm scores (calculation of ANI based on the MUMmer algorithm which is more adapted to compare genomes with high degree of similarity (51, 52)).

### HiSST assignation

The HiSST identification of an isolate is performed with a R script, and new HiSST profiles for unknown isolates are added to the HiSST database using the same script. The HiSST scheme database and the R script called “HiSST-Assignation” are available on GitHub at URL: https://github.com/TBourd/R_scripts_for_HiSST_scheme.

### Accession number(s)

Raw sequencing reads have been deposited in the Sequence Read Archive of the NCBI in the BioProject <u>PRJNA729113</u>. Isolates raw sequencing reads are in BioSamples SAMN19117128 to SAMN19117139, and eDNA raw sequencing reads are in BioSamples SAMN19110658 to SAMN19110711. Assembled genomes of the environmental strain BD1b-2wD and clinical strains ED3957, ED3958, ED3959 have been deposited in the same BioProject <u>PRJNA729113</u> into BioSamples accessions SAMN19232018 to SAMN19232021.

## Results and Discussion

### Design of the HiSST scheme

Thirty two out of the 3,301 genes of *S. marcescens* pangenome were identified as the most variable with 29 to 30 alleles per gene. Only the most specific and discriminatory genes were kept, after stepwise alignment, examination of polymorphism amongst *S. marcescens* genomes and specificity check (Fig. 1). This led to the selection of 15 loci for subsequent analyses listed in Table S4. The minimal number of loci included in the HiSST scheme was selected by topology data analysis of concatenated HiSST loci and genome similarity (Fig. 2). We found that the dendrogram built from three loci has a similar topology to the cognate clustering analysis comprising the 15 most discriminatory loci, according to the downward loci selection procedure. This led to the selection of the three loci located in genes *gabR, bssA* and *dhaM* for the HiSST scheme. Each locus was discriminant, with more than 20% of nucleotide dissimilarity between *S. marcescens* strains and other species (Fig. S1). The locus *gabR* is more specific to *S. marcescens* species than *bssA*, followed by *dhaM*, with respectively more than 26%, 17% and 14% of nucleotide dissimilarity with *S. ficaria*, the closest relative species of *S. marcescens* for these loci. The HiSST scheme based on these 3 loci differentiates most *S. marcescens* strains better than the ANI score based on their whole genomes (Fig. 3). The pairwise ANIb genome similarity score is over 94% in its ability to distinguish *S. marcescens* strains from other *Serratia* species while only a few strains of *Serratia* spp. have more than 70% nucleotide identity with the three selected loci of *S. marcescens*. Classification of *S. marcescens* strains based on the HiSST-scheme is congruent with classification scheme relying on complete genome sequences (Fig. 2). A few differences between HiSST and whole genome-based classification were noticed amongst strains sharing more than 99.9% ANI score. At such a high similarity level, threshold delineating species, strains or clones is empirical, depending on examined species. For example, *P. aeruginosa* has high genomic plasticity mainly due to frequent horizontal gene transfers (53, 54), while *S. marcescens* has a higher genetic diversity at the sequence level according to PGAdb-builder results, with also genome flexibility (55). Additional factors to consider include the study context (e.g. precautionary principle for epidemiological studies tend to identify highly similar but not identical strains as non-clonal strain) and the method used (i.e. depending on the sensitivity of the molecular typing method and the sequencing platform used, the evolution of the technology and knowledge). As a whole, the minimal similarity threshold amongst *S. marcescens* strains is 89% for the three loci (Fig. S2 and based on BLAST results).

**Figure 1:**
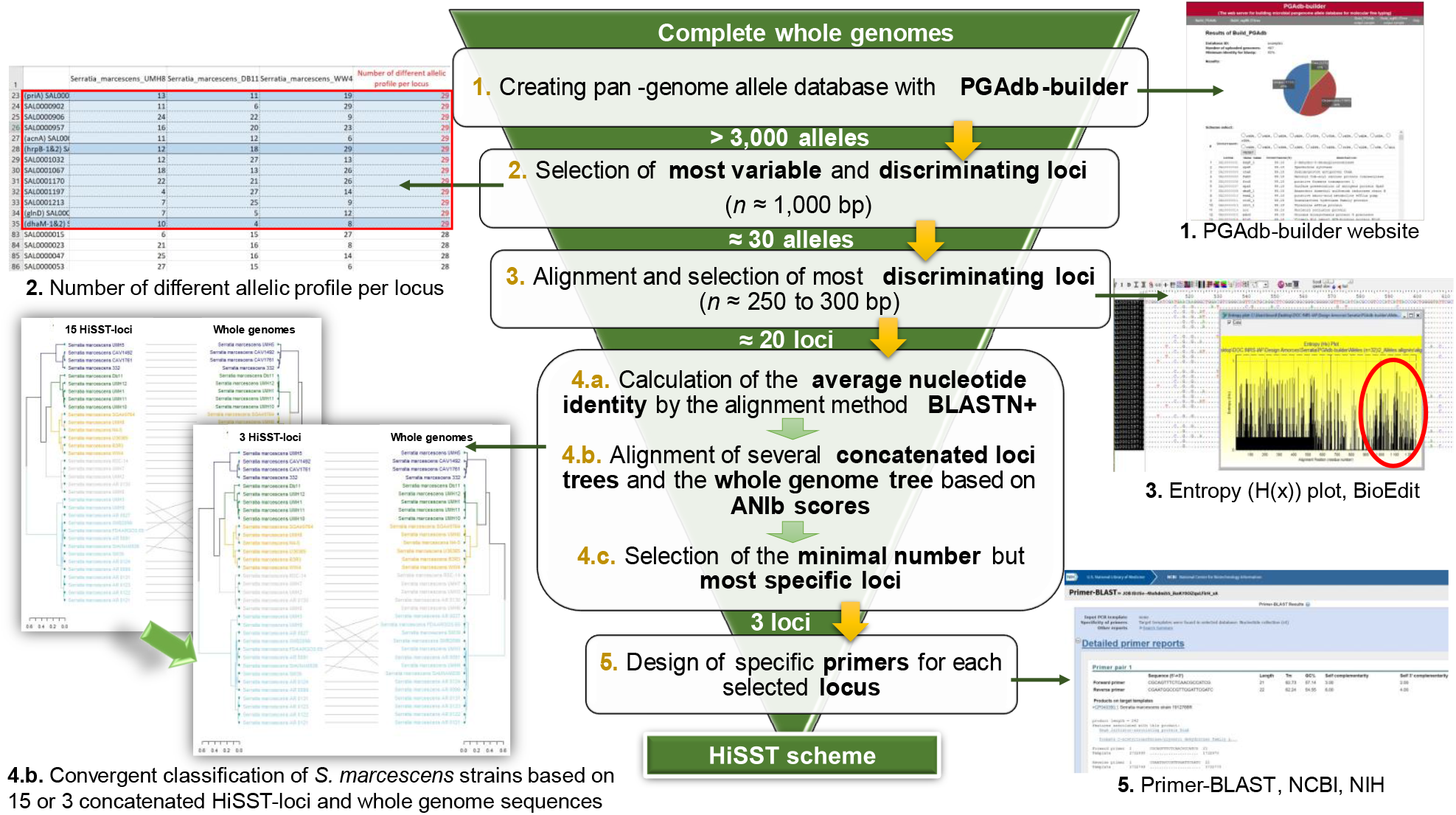
Step-by-step approach of the method used to develop the HiSST scheme of *S. marcescens.*

**Figure 2:**
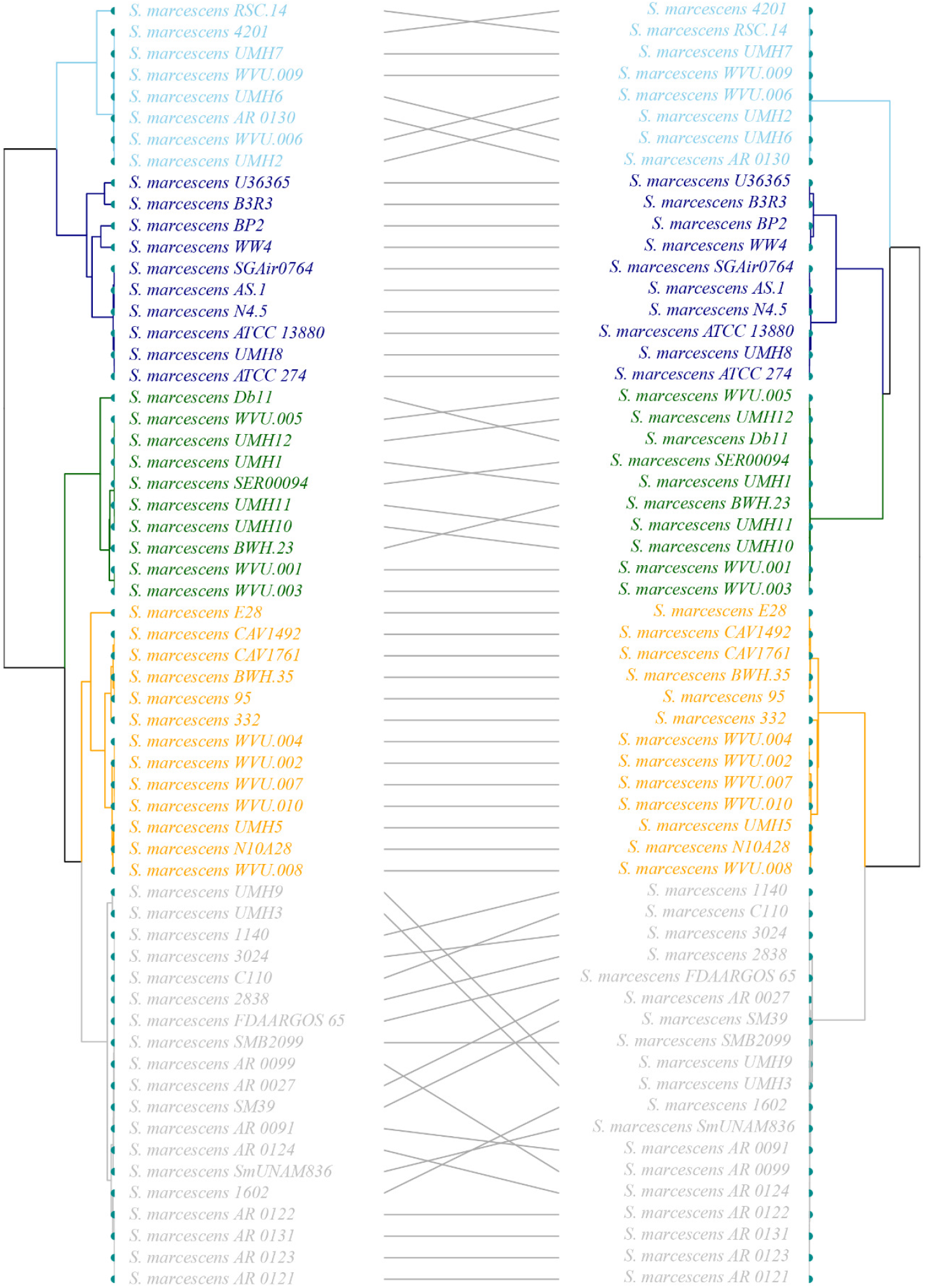
Convergent classification of *S. marcescens* strains based on the HiSST scheme and whole genome sequences. UPGMA dendrogram based on the ANIb score of concatenated loci selected for (A) HiSST scheme and (B) genome similarity to discriminate strains of *Serratia marcescens*.

**Figure 3:**
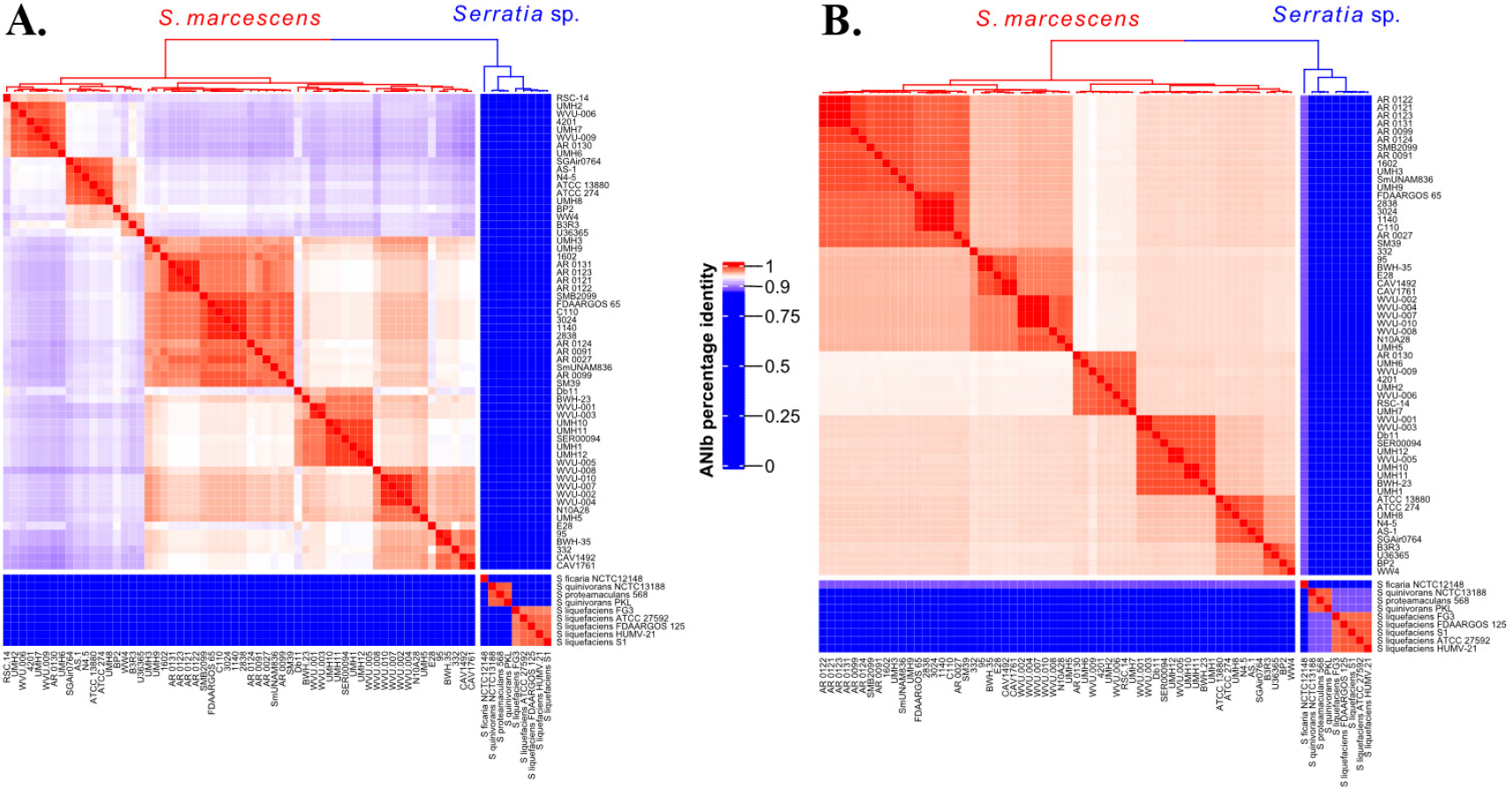
Discrimination of *Serratia* spp. based on the HiSST scheme and whole genome sequences. The heat-map reports the ANIb score of (A) the three concatenated loci of the HiSST scheme and (B) genome similarity. *S. ficaria* (n=1), *S. quinivorans* (n=2), *S. proteamaculans* (n=1) and *S. liquefaciens* (n=5) were included as outgroup.

### Validation and application of the HiSST scheme

Specificity of primers targeting *gabR, bssA* and *dhaM* loci was first confirmed by Blast searches against RefSeq non-redundant proteins database. The efficacy and the specificity of the PCR assays was further confirmed with reference strains (Fig. S3). PCR amplicon of the correct size was observed for all *S. marcescens* strains (*n* = 15) but not for *Serratia* sp. and other species of gammaproteobacteria. An accuracy test of HiSST scheme, including the bioinformatic procedure utilized to assign alleles to ST, was realized with reference strains *S. marcescens* Db11 and Db10. Genomic DNA of both strains was subjected to PCR amplicon sequencing with an average allocation of 1,000 reads per library. The HiSST-profile (ST 1) of both strains corresponded to the expected profile with a single ASV for each gene, supporting the accuracy of HiSST procedure and parameters utilized in sequence quality control (Fig. 4).

**Figure 4:**
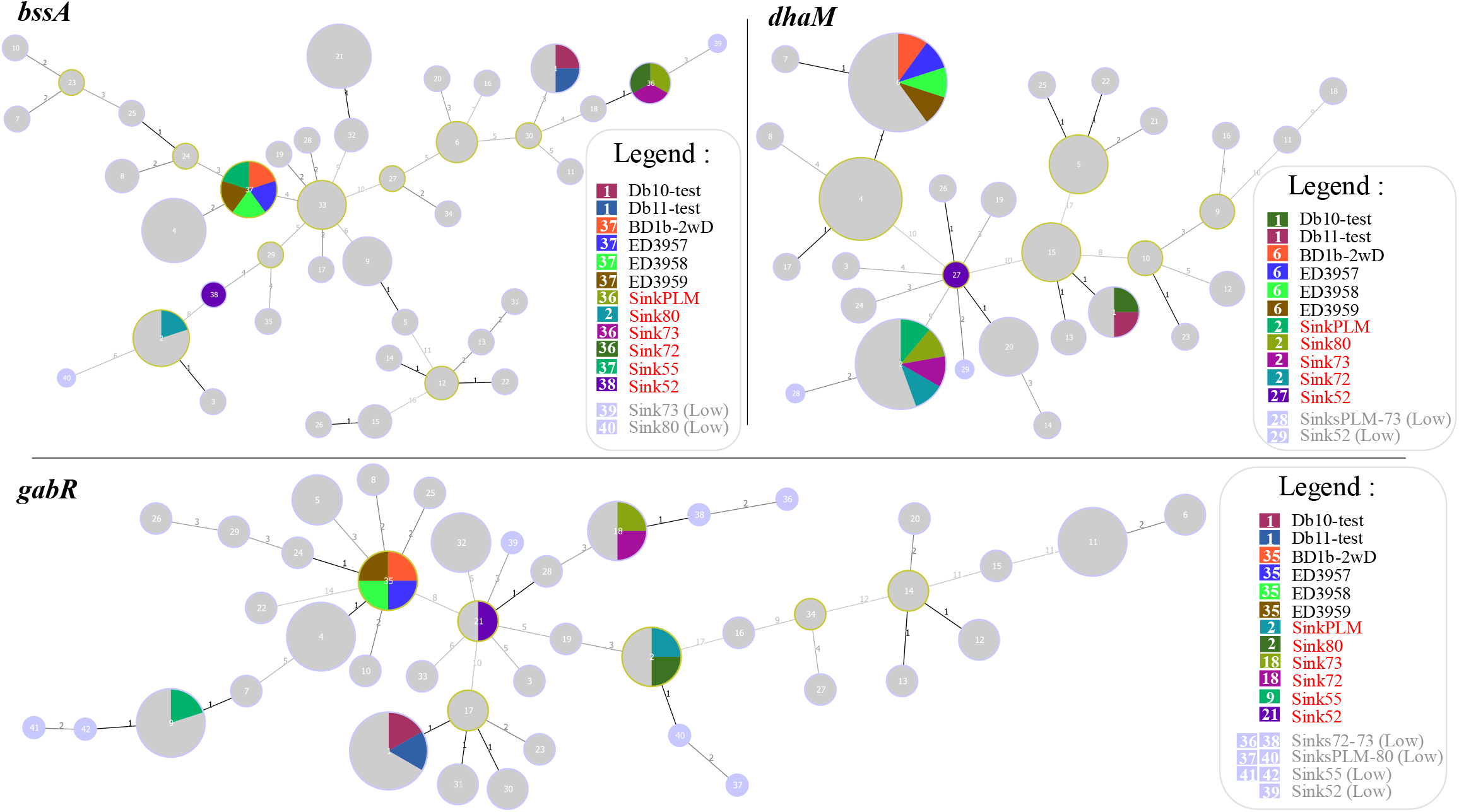
Minimum spanning trees based on SNP analysis of *S. marcescens* and eDNA, using *S. marcescens* Db10 as a reference. The distance labels represent the number of discriminating SNPs between neighbouring genotypes. Each pie chart label refers to ST identifier of the corresponding locus. Reference genomes are represented in grey and isolates or sampled sinks are represented by the colour legend in pie charts. Dominant STs of eDNA are represented with red font characters in the legend box whereas grey characters correspond to ST of eDNA in low abundance.

The method was applied to two different case studies realized in the same NICU. The first case study sought to compare the ST profile of a strain isolated from the sink-drain environment (BD1b-2wD) with three clinical strains (ED3657, ED3658, ED3659) from patients admitted in that NICU, where an outbreak occurred – as determined by the Infection prevention and control team based on PFGE profiles, relatedness in space and time. Molecular typing of the environmental strain BD1b-2wD and clinical strains revealed very close relatedness, all four having an identical HiSST-profile ST 47 (Fig. 5A). WGS was done for each strain to challenge HiSST scheme result. Pairwise comparison of contiguated genomes confirmed the high degree of similarity between each strain (ANIm > 99.7%). In principle, *bssA* and *gabR* are sufficient to ensure diversity coverage of ST represented in genome database (Fig. 5B) but inclusion of *dhaM* in the HiSST-scheme is included to prevent false-negative results (i.e., in the case where the targeted gene is absent or subject to unknown mutations) and allows to distinguish environmental or clinical origin of *S. marcescens* strains for culture-based diagnostic (Fig. 5A). These results suggest that the environmental BD1b-2wD strain and clinical isolates descend from a single cell, while providing supplementary experimental evidence supporting the specificity of the HiSST scheme.

**Figure 5:**
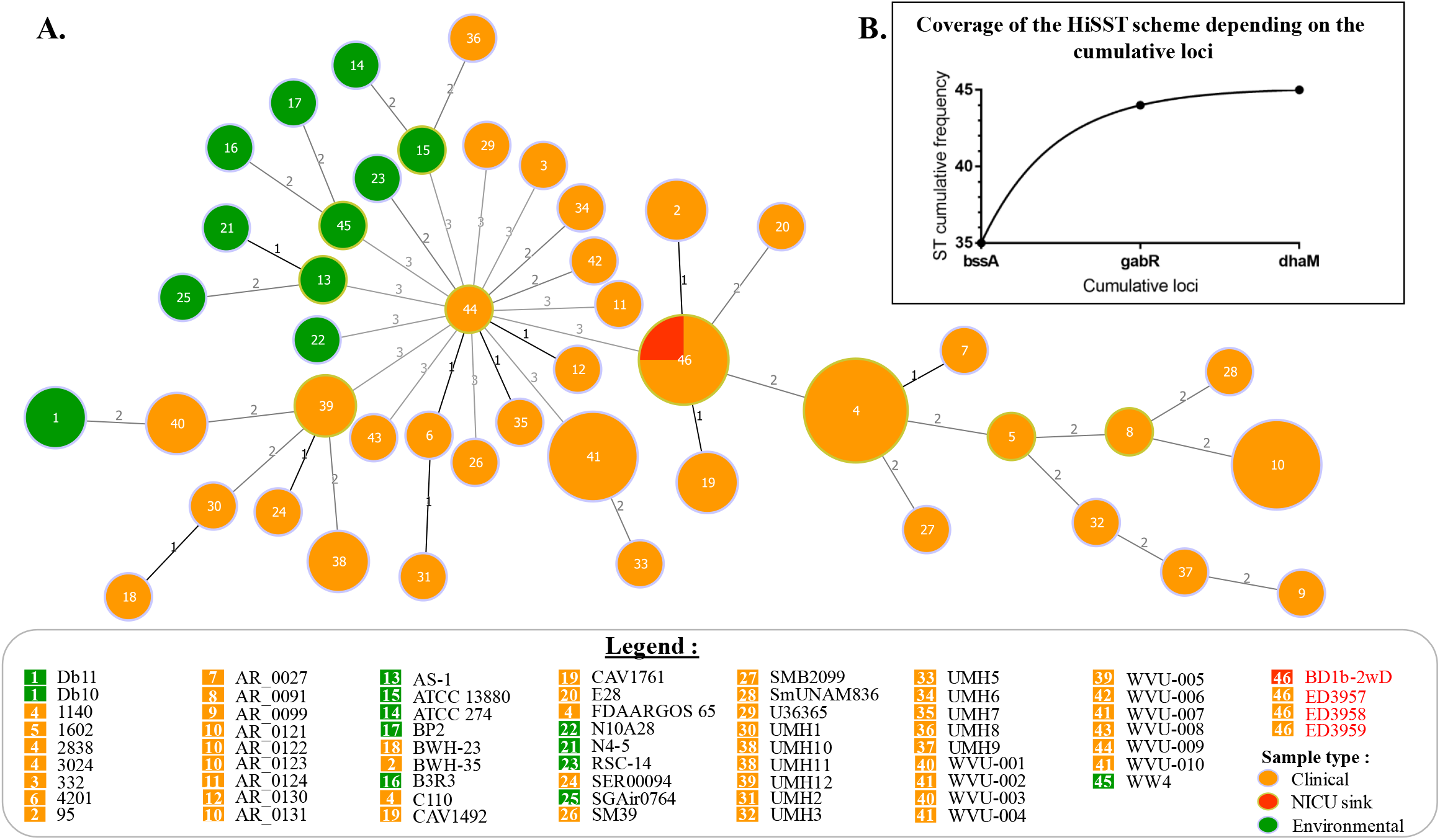
Relationship amongst the ST profile of reference strains and isolates and diversity coverage of the HiSST scheme. (A) A minimum spanning tree based on MLST analysis of HiSST scheme is represented with distance labels corresponding to the number of discriminating alleles and pie chart labels referring to the ST identifier of the HiSST scheme. Orange nodes correspond to clinical isolates, the red node to the isolate from NICU sink-drain, and the green nodes to environmental isolates. In the legend box, strains represented by red font characters correspond to unknown clinical (ED3957, ED3958, ED3959) and sink-drain (BD1b-2wD) isolates from this study. (B) Cumulative frequency of ST depending on the number of loci included in the HiSST scheme.

The second case study was conducted to explore diversity of *S. marcescens* by applying the HiSST method to environmental DNA (eDNA). PCR amplicon sequencing of each loci was done to report diversity of each ST-locus separately for a culture-independent epidemiological investigation. All retrieved ASV sequences were specific to *S. marcescens*. Diversity amongst the 19 sinks was modest, with 11, 6 and 4 different alleles of *gabR, bssA* and *dhaM* found, respectively (Fig. 4). A single allele was dominant in each sample, with a relative abundance of 70-100% (Table S2). Either a single or two allele(s) per sample were observed for *gabR* and *dhaM* loci, whereas *gabR* was represented by up to three different alleles per sample. For the three HiSST loci, rare alleles differ from the dominant allele in the same sample by 1-6 SNPs, suggesting the presence of other strains in the drain. Artificial inflation of diversity caused by sequencing errors is less likely due to the stringent filtering process of sequences (cf. Materials and methods) and the low error probability of incorrect base-call for short sequences (56). The intercomparison of ST profiles amongst the 19 sinks of the NICU was done to infer potential epidemiological links (Fig. 6). The most straightforward link between sink environments is the case where ST profiles are identical. This situation was observed in sinks #72 and #73 for dominants ASV (*bssA*-ST 36, *gabR*-ST 18, *dhaM*-ST 2) that are likely colonized by the same *S. marcescens* strain. This link is supported by the proximity of both sinks in the NICU, with the same drain connection and interconnection through handwashing (57, 58). The sink #PLM shared two ST detected in sinks #72 and #73 (*bssA*-ST 36 and *dhaM*-ST 2) and two ST in sink #80 (*gabR*-ST 2 and *dhaM*-ST 2). This result suggests an epidemiological link between the four sinks related to one another by the sink #PLM (that is used for the initial handwashing at the NICU entrance). Finally, HiSST-profile (ST 2) of sink #80 is identical to *S. marcescens* 95 and BWH-35 strains included in the reference genome database, suggesting the colonization by a taxonomically closely-related strain. *S. marcescens* 95 and BWH-35 were isolated from sputum in a Boston hospital (USA) and are most likely variants of the same strain.

**Figure 6:**
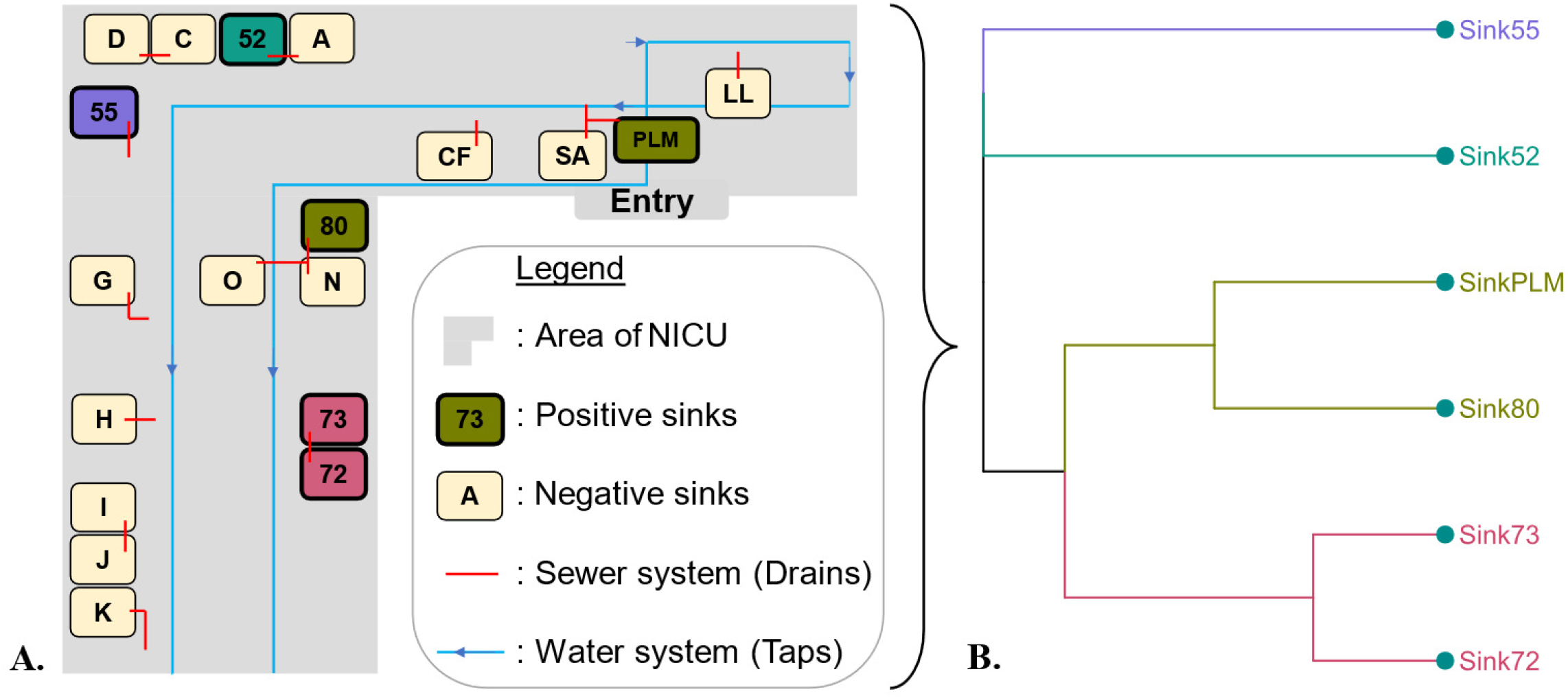
Survey of *Serratia marcescens* in sink drains of a NICU. (A) A Scheme of the surveyed NICU is depicted along an (B) UPGMA dendrogram based on Jaccard distance computed with the HiSST profile of *gabR, bssA and dhaM* loci amongst sink drains that showed positive PCR amplifications.

A limitation of the method was noticed in sink #55 where no PCR detection of *dhaM* was observed with positive amplification of *gabR* and *bssA* genes. Although this can be explained by low level of *S. marcescens* in this sink combined with different amplification efficiencies between the three reactions, examination of future genome sequences deposited in public database will be necessary to confirm the ubiquitous distribution of *dhaM* in *S. marcescens*.

These case studies illustrate the strengths of the HiSST scheme to identify clones and its broad applicability for epidemiological investigations. Beyond the conventional application of the method to genotype isolates, examination of eDNA offers a complementary tool for the source tracking of opportunistic pathogens. This could be done by the monitoring of bacterial succession in NICU environment and patient samples through HiSST eDNA profiling. Under that framework, a convergence of HiSST profiles along spatial or temporal sampling sequences would provide strong evidence of opportunistic pathogen transfer across different environments.

In contrast to conventional application for isolate identification, HiSST profile analysis from eDNA is less prone to misinterpretation or aborted analysis for samples displaying no signal for certain genes. Indeed, the pairwise comparison of HiSST bacterial profiles can be expressed as a pairwise Jaccard distance computed with presence or absence score for detected or non-detected ST, respectively. Downstream clustering and multivariate analyses offer options to correlate ST distribution patterns with environmental features encompassing sink location, utilization, and microbial diversity (Fig. 6). Although this approach is a gold standard in microbial ecology, the second case study presented in this article is the first culture-independent application of ST profile analysis of opportunistic pathogens for epidemiologic survey.

In conclusion, a combination of *in silico* analyses led to the development of a powerful HiSST assay to identify isolates of *S. marcescens* species. The approach relying on pangenome examination rather than selection of conventional housekeeping genes contributed to the method specificity. For instance, conventional MLST schemes for *P. aeruginosa* and *S. maltophilia* are less specific than the HiSST method developed here for *S. marcescens*. Application of the procedure presented in this article to these other opportunistic pathogens of environmental origin led to more robust HiSST-schemes (T. Bourdin et al., unpublished). Despite the precision of the method presented here, specificity and coverage of the HiSST scheme will require regular validation and update with the addition of new genome sequences in public databases. The bioinformatic pipeline implemented here or alternative methods (59) will facilitate regular update of the HiSST scheme. This fact holds true for any molecular classification tool. Even though comparison of whole genomes appears as the most robust method (12), public genome databases contain contaminations that may introduce biases for the identification of highly similar strains (60). In addition, the high proportion of similar or identical genes in whole genome hides some dissimilarities between isolates, while HiSST highlights the most discriminating alleles. Thus, a combination of whole genome sequencing and high discriminatory molecular typing method is recommended for culture-dependant epidemiological investigation (61). Beside isolate identification, the HiSST method proved efficient for ST comparison and source tracking purposes of *S. marcescens* in eDNA samples without the need for culture.

Based on these results, the following epidemiological interpretations for molecular typing of isolates when using HiSST scheme are proposed: (i) isolates that are identified by at least 2 of 3 HiSST-loci are confirmed as *S. marcescens*, (ii) isolates with an identical HiSST-profile (i.e. identical *gabR*-ST, *bssA*-ST and *dhaM*-ST) are most likely clones and belong to the same genotype, (iii) isolates that differ by 2 or 3 HiSST-loci are mostly unrelated and do not belong to the same genotype. For an epidemiological survey on eDNA samples when using HiSST scheme described here, the following interpretations are proposed: (i) eDNA samples with ST corresponding to the HiSST scheme indicate the presence of *S. marcescens*, (ii) eDNA samples with several ST of one HiSST-locus indicate the presence of several *S. marcescens* strains, and (iii) samples with identical HiSST-profile are harbouring by very closely related strains and sampled environment are most likely linked.

## Acknowledgments

We thank the hospital staff for help in sampling, Ann Brassinga (Department of Microbiology, University of Manitoba), Jonathan J. Ewbank (Centre d’Immunologie de Marseille-Luminy, Aix-Marseille University, Marseille, France), Sabine Favre-Bonté (Université Lyon 1, UMR CNRS 5557 Ecologie Microbienne, Lyon, France), and the Laboratoire de santé publique du Québec for providing reference strains.

This work was supported by NSERC and CIHR through the IRC Industrial Chair on Drinking Water and the Collaborative Health Research Program funding (CHRP 523790-18).

## Supplementary figures

**Figure S1:**
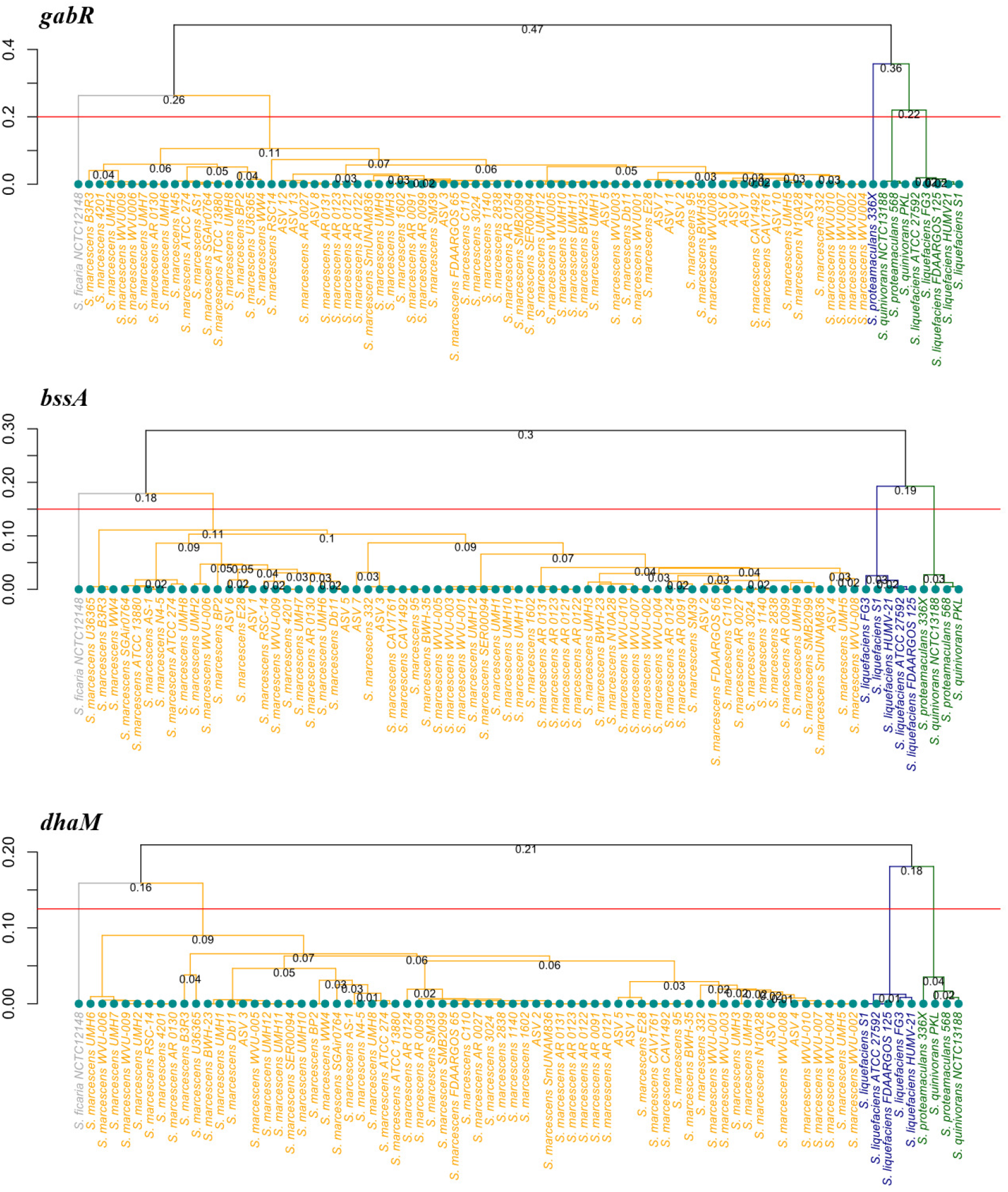
UPGMA trees of each locus selected for HiSST scheme between *Serratia sp*. strains and environmental ASV, based on Jukes-Cantor distance. Each cluster gathers strains with more than 90% of similarity.

**Figure S2:**
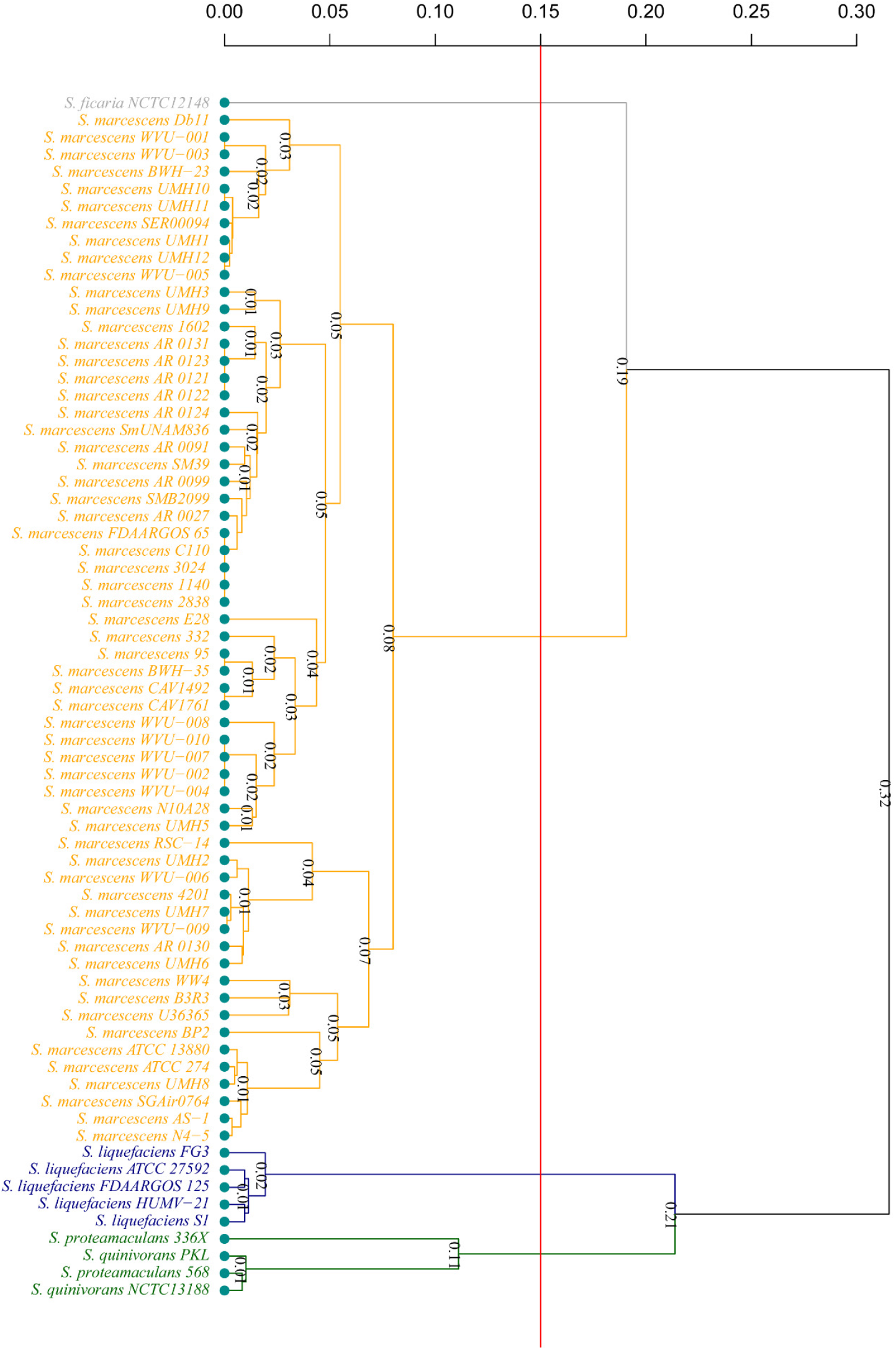
UPGMA trees for topological data analysis of concatenated loci selected for HiSST scheme based on ANIb score between *Serratia sp*. strains. Each cluster gathers strains with more than 85% of nucleotide similarity.

**Figure S3:**
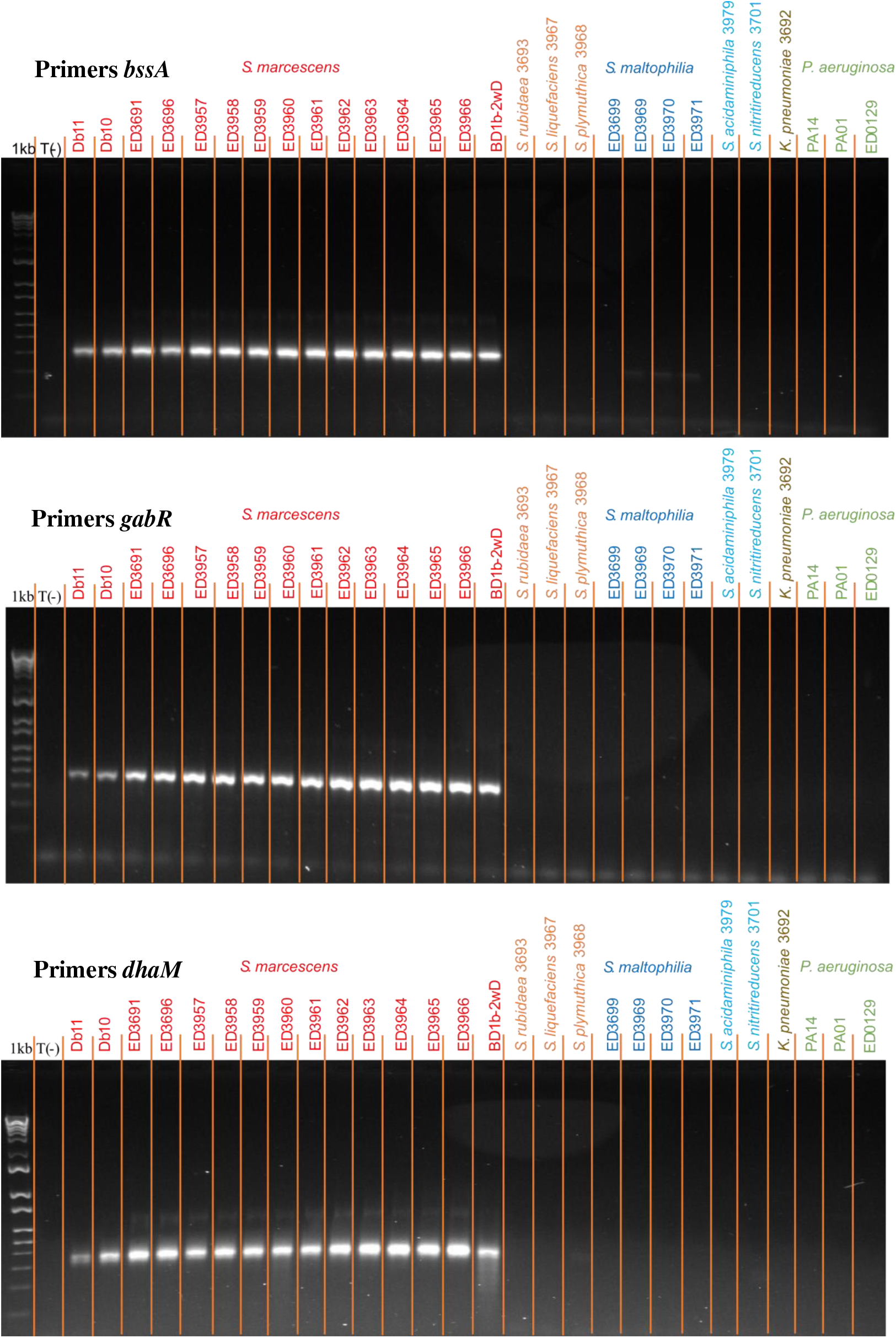
Tests and validation *in-vitro* of primers designed for HiSST scheme.

## References

1. Regev-Yochay G, Smollan G, Tal I, Pinas Zade N, Haviv Y, Nudelman V, Gal-Mor O, Jaber H, Zimlichman E, Keller N, Rahav G. 2018. Sink traps as the source of transmission of OXA-48-producing Serratia marcescens in an intensive care unit. Infect Control Hosp Epidemiol 39:1307–1315.

2. Bédard E, Laferrière C, Charron D, Lalancette C, Renaud C, Desmarais N, Déziel E, Prévost M. 2015. Post-outbreak investigation of Pseudomonas aeruginosa faucet contamination by quantitative polymerase chain reaction and environmental factors affecting positivity. Infect Control Hosp Epidemiol 36:1337–1343.

3. Lalancette C, Charron D, Laferrière C, Dolcé P, Déziel E, Prévost M, Bédard E. 2017. Hospital Drains as Reservoirs of Pseudomonas aeruginosa: multiple-locus variable-number of tandem repeats analysis genotypes recovered from faucets, sink surfaces and patients. 3. Pathogens 6:36.

4. Inouye M, Conway TC, Zobel J, Holt KE. 2012. Short read sequence typing (SRST): multi-locus sequence types from short reads. BMC Genomics 13:338.

5. Boers SA, Reijden WA van der, Jansen R. 2012. High-Throughput Multilocus Sequence Typing: Bringing Molecular Typing to the Next Level. PLOS ONE 7:e39630.

6. Sabat AJ, Budimir A, Nashev D, Sá-Leão R, Dijl JM van, Laurent F, Grundmann H, Friedrich AW, on behalf of the ESCMID Study Group of Epidemiological Markers (ESGEM). 2013. Overview of molecular typing methods for outbreak detection and epidemiological surveillance. Eurosurveillance 18:20380.

7. Basset P, Blanc DS. 2014. Fast and simple epidemiological typing of Pseudomonas aeruginosa using the double-locus sequence typing (DLST) method. Eur J Clin Microbiol Infect Dis 6.

8. de Been M, Pinholt M, Top J, Bletz S, Mellmann A, van Schaik W, Brouwer E, Rogers M, Kraat Y, Bonten M, Corander J, Westh H, Harmsen D, Willems RJL. 2015. Core genome multilocus sequence typing scheme for high-resolution typing of Enterococcus faecium. Journal of Clinical Microbiology 53:3788–3797.

9. Chen Y, Frazzitta AE, Litvintseva AP, Fang C, Mitchell TG, Springer DJ, Ding Y, Yuan G, Perfect JR. 2015. Next generation multilocus sequence typing (NGMLST) and the analytical software program MLSTEZ enable efficient, cost-effective, high-throughput, multilocus sequencing typing. Fungal Genetics and Biology 75:64–71.

10. Tewolde R, Dallman T, Schaefer U, Sheppard CL, Ashton P, Pichon B, Ellington M, Swift C, Green J, Underwood A. 2016. MOST: a modified MLST typing tool based on short read sequencing. PeerJ 4:e2308.

11. Maiden MC, Bygraves JA, Feil E, Morelli G, Russell JE, Urwin R, Zhang Q, Zhou J, Zurth K, Caugant DA, others. 1998. Multilocus sequence typing: a portable approach to the identification of clones within populations of pathogenic microorganisms. Proceedings of the National Academy of Sciences 95:3140–3145.

12. Maiden MCJ, Jansen van Rensburg MJ, Bray JE, Earle SG, Ford SA, Jolley KA, McCarthy ND. 2013. MLST revisited: the gene-by-gene approach to bacterial genomics. Nat Rev Microbiol 11:728–736.

13. Bleidorn C, Gerth M. 2018. A critical re-evaluation of multilocus sequence typing (MLST) efforts in Wolbachia. FEMS Microbiology Ecology 94.

14. Grimont PAD, Grimont F. 1978. The Genus Serratia. Annu Rev Microbiol 32:221–248.

15. Hejazi A, Falkiner FR. 1997. Serratia marcescens. Journal of Medical Microbiology 46:903–912.

16. Villari P, Crispino M, Salvadori A, Scarcella A. 2001. Molecular epidemiology of an outbreak of Serratia marcescens in a neonatal intensive care unit. Infect Control Hosp Epidemiol 22:630–634.

17. Assadian O, Berger A, Aspöck C, Mustafa S, Kohlhauser C, Hirschl AM. 2002. Nosocomial outbreak of Serratia marcescens in a neonatal intensive care unit. Infect Control Hosp Epidemiol 23:457–461.

18. Milisavljevic V, Wu F, Larson E, Rubenstein D, Ross B, Drusin LM, Della-Latta P, Saiman L. 2004. Molecular epidemiology of Serratia marcescens outbreaks in two neonatal intensive care units. Infect Control Hosp Epidemiol 25:719–722.

19. Maragakis LL, Winkler A, Tucker MG, Cosgrove SE, Ross T, Lawson E, Carroll KC, Perl TM. 2008. Outbreak of multidrug-resistant Serratia marcescens infection in a neonatal intensive care unit. Infect Control Hosp Epidemiol 29:418–423.

20. Zingg W, Soulake I, Baud D, Huttner B, Pfister R, Renzi G, Pittet D, Schrenzel J, Francois P. 2017. Management and investigation of a Serratia marcescens outbreak in a neonatal unit in Switzerland – the role of hand hygiene and whole genome sequencing. Antimicrobial Resistance & Infection Control 6:125.

21. Åttman E, Korhonen P, Tammela O, Vuento R, Aittoniemi J, Syrjänen J, Mattila E, Österblad M, Huttunen R. 2018. A Serratia marcescens outbreak in a neonatal intensive care unit was successfully managed by rapid hospital hygiene interventions and screening. Acta Paediatrica 107:425–429.

22. Martineau C, Li X, Lalancette C, Perreault T, Fournier E, Tremblay J, Gonzales M, Yergeau É, Quach C. 2018. Serratia marcescens outbreak in a neonatal intensive care unit: new insights from next-generation sequencing applications. Journal of Clinical Microbiology 56:235–18.

23. Moles L, Gómez M, Moroder E, Jiménez E, Escuder D, Bustos G, Melgar A, Villa J, del Campo R, Chaves F, Rodríguez JM. 2019. Serratia marcescens colonization in preterm neonates during their neonatal intensive care unit stay. Antimicrobial Resistance & Infection Control 8:135.

24. Cristina ML, Sartini M, Spagnolo AM. 2019. Serratia marcescens infections in neonatal intensive care units (NICUs). 4. International Journal of Environmental Research and Public Health 16:610.

25. Varsha G, Shiwani S, Kritika P, Poonam G, Deepak A, Jagdish C. 2021. Serratia no longer an opportunistic uncommon pathogen – case series & review of literature. Infectious Disorders -Drug Targets 21:1–1.

26. Johnson J, Quach C. 2017. Outbreaks in the neonatal ICU: a review of the literature. Current Opinion in Infectious Diseases 30:395–403.

27. Friedman ND, Kotsanas D, Brett J, Billah B, Korman TM. 2008. Investigation of an outbreak of Serratia marcescens in a neonatal unit via a case-control study and molecular typing. American Journal of Infection Control 36:22–28.

28. Rossen JWA, Dombrecht J, Vanfleteren D, Bruyne KD, Belkum A van, Rosema S, Lokate M, Bathoorn E, Reuter S, Grundmann H, Ertel J, Higgins PG, Seifert H. 2019. Epidemiological typing of Serratia marcescens isolates by whole-genome multilocus sequence typing. Journal of Clinical Microbiology 57:1652–1618.

29. Liu Y-Y, Chiou C-S, Chen C-C. 2016. PGAdb-builder: A web service tool for creating pan-genome allele database for molecular fine typing. Scientific Reports 6:36213.

30. Hall, T.A. 1999. BioEdit: a user-friendly biological sequence alignment editor and analysis program for Windows 95/98/NT. Nucl Acids Symp Ser 95–98.

31. Figueras MJ, Beaz-Hidalgo R, Hossain MJ, Liles MR. 2014. Taxonomic affiliation of new genomes should be verified using average nucleotide identity and multilocus phylogenetic analysis. Genome Announc 2:927–914.

32. Pritchard L, Glover RH, Humphris S, Elphinstone JG, Toth IK. 2015. Genomics and taxonomy in diagnostics for food security: soft-rotting enterobacterial plant pathogens. Anal Methods 8:12–24.

33. R Core Team. 2021. R: A language and environment for statistical computing. R Foundation for Statistical Computing, Vienna, Austria.

34. Suzuki R, Shimodaira H. 2006. Pvclust: an R package for assessing the uncertainty in hierarchical clustering. Bioinformatics 22:1540–1542.

35. Galili T. 2015. dendextend: an R package for visualizing, adjusting and comparing trees of hierarchical clustering. Bioinformatics 31:3718–3720.

36. Wickham H, Averick M, Bryan J, Chang W, McGowan LD, François R, Grolemund G, Hayes A, Henry L, Hester J, Kuhn M, Pedersen TL, Miller E, Bache SM, Müller K, Ooms J, Robinson D, Seidel DP, Spinu V, Takahashi K, Vaughan D, Wilke C, Woo K, Yutani H. 2019. Welcome to the Tidyverse. Journal of Open Source Software 4:1686.

37. Gu Z, Gu L, Eils R, Schlesner M, Brors B. 2014. circlize implements and enhances circular visualization in R. Bioinformatics 30:2811–2812.

38. Gu Z, Eils R, Schlesner M. 2016. Complex heatmaps reveal patterns and correlations in multidimensional genomic data. Bioinformatics 32:2847–2849.

39. Ye J, Coulouris G, Zaretskaya I, Cutcutache I, Rozen S, Madden TL. 2012. Primer-BLAST: A tool to design target-specific primers for polymerase chain reaction. BMC Bioinformatics 13:134.

40. Berkowitz DM, Lee WS. 1973. A selective medium for isolation and identification of Serratia marcescens. Abstracts of the Annual Meeting of the American Society for Microbiology 1973 105.

41. Durand A-A, Bergeron A, Constant P, Buffet J-P, Déziel E, Guertin C. 2015. Surveying the endomicrobiome and ectomicrobiome of bark beetles: The case of Dendroctonus simplex. 1. Scientific Reports 5:17190.

42. Martin M. 2011. Cutadapt removes adapter sequences from high-throughput sequencing reads. 1. EMBnet.journal 17:10–12.

43. Callahan BJ, McMurdie PJ, Rosen MJ, Han AW, Johnson AJA, Holmes SP. 2016. DADA2: High-resolution sample inference from Illumina amplicon data. 7. Nature Methods 13:581– 583.

44. Morgan M, Lawrence M, Anders S. 2021. ShortRead: FASTQ input and manipulation. Bioconductor version: Release (3.12).

45. Pagès H, Aboyoun P, Gentleman R, DebRoy S. 2021. Biostrings: Efficient manipulation of biological strings. Bioconductor version: Release (3.12).

46. Nascimento M, Sousa A, Ramirez M, Francisco AP, Carriço JA, Vaz C. 2017. PHYLOViZ 2.0: providing scalable data integration and visualization for multiple phylogenetic inference methods. Bioinformatics 33:128–129.

47. Bolger AM, Lohse M, Usadel B. 2014. Trimmomatic: a flexible trimmer for Illumina sequence data. Bioinformatics 30:2114–2120.

48. Prjibelski A, Antipov D, Meleshko D, Lapidus A, Korobeynikov A. 2020. Using SPAdes De Novo Assembler. Current Protocols in Bioinformatics 70:e102.

49. Wick RR, Schultz MB, Zobel J, Holt KE. 2015. Bandage: interactive visualization of de novo genome assemblies. Bioinformatics 31:3350–3352.

50. Assefa S, Keane TM, Otto TD, Newbold C, Berriman M. 2009. ABACAS: algorithm-based automatic contiguation of assembled sequences. Bioinformatics 25:1968–1969.

51. Richter M, Rosselló-Móra R. 2009. Shifting the genomic gold standard for the prokaryotic species definition. PNAS 106:19126–19131.

52. Kurtz S, Phillippy A, Delcher AL, Smoot M, Shumway M, Antonescu C, Salzberg SL. 2004. Versatile and open software for comparing large genomes. Genome Biol 5:R12.

53. Diggle SP, Whiteley M. 2020. Microbe Profile: Pseudomonas aeruginosa: opportunistic pathogen and lab rat. Microbiology (Reading) 166:30–33.

54. Freschi L, Vincent AT, Jeukens J, Emond-Rheault J-G, Kukavica-Ibrulj I, Dupont M-J, Charette SJ, Boyle B, Levesque RC. 2019. The Pseudomonas aeruginosa pan-genome provides new insights on its population structure, horizontal gene transfer, and pathogenicity. Genome. Biology and Evolution 11:109–120.

55. Iguchi A, Nagaya Y, Pradel E, Ooka T, Ogura Y, Katsura K, Kurokawa K, Oshima K, Hattori M, Parkhill J, Sebaihia M, Coulthurst SJ, Gotoh N, Thomson NR, Ewbank JJ, Hayashi T. 2014. Genome evolution and plasticity of Serratia marcescens, an important multidrug-resistant nosocomial pathogen. Genome Biol Evol 6:2096–2110.

56. Ewing B, Green P. 1998. Base-calling of automated sequencer traces using Phred. II. Error Probabilities. Genome Res 8:186–194.

57. Franco LC, Tanner W, Ganim C, Davy T, Edwards J, Donlan R. 2020. A microbiological survey of handwashing sinks in the hospital built environment reveals differences in patient room and healthcare personnel sinks. 1. Scientific Reports 10:8234.

58. Wingender J. 2011. Hygienically relevant microorganisms in biofilms of man-made water systems, p. 189–238. In Flemming, H-C, Wingender, J, Szewzyk, U (eds.), Biofilm Highlights. Springer, Berlin, Heidelberg.

59. Gaiarsa S, Batisti Biffignandi G, Esposito EP, Castelli M, Jolley KA, Brisse S, Sassera D, Zarrilli R. 2019. Comparative analysis of the two Acinetobacter baumannii multilocus sequence typing (MLST) schemes. Front Microbiol 10:930.

60. Steinegger M, Salzberg SL. 2020. Terminating contamination: large-scale search identifies more than 2,000,000 contaminated entries in GenBank. Genome Biology 21:115.

61. Magalhães B, Valot B, Abdelbary MMH, Prod’hom G, Greub G, Senn L, Blanc DS. 2020. Combining standard molecular typing and whole genome sequencing to investigate Pseudomonas aeruginosa epidemiology in intensive care units. Front Public Health 8:3.

